# Whole-brain protein profiling using organ-scale multiplexed immunolabeling and image co-registration

**DOI:** 10.64898/2026.05.06.723275

**Authors:** Sehun Kim, Hyeongryool Park, Wonjin Cho, Sangbin Yoo, Thananya Charoenpattarawut, Christopher E. Pearson, Young-Gyun Park

**Affiliations:** Department of Bio and Brain Engineering, Korea Advanced Institute of Science and Technology (KAIST), Daejeon, Republic of Korea; Brain and Cognitive Engineering Program, Korea Advanced Institute of Science and Technology (KAIST), Daejeon, Republic of Korea; Department of Biological Sciences, Korea Advanced Institute of Science and Technology (KAIST), Daejeon, Republic of Korea; KAIST Institute for Health Science and Technology, Korea Advanced Institute of Science and Technology (KAIST), Daejeon, Republic of Korea; Genetics & Genome Biology, The Hospital for Sick Children, Toronto, ON, Canada; Department of Molecular Genetics, University of Toronto, Toronto, ON, Canada

## Abstract

Proteins are major drivers of biological functions. Single-cell, organ-scale multiplexed protein imaging can reveal high-dimensional molecular and structural features of individual cells and their interactions, enabling an in-depth understanding of complex biological systems. However, achieving such imaging has remained elusive due to hurdles in multiplexed immunolabeling (mIF) of intact organs and integrative image analysis. Here, we present 3D CYCLIC, an organ-scale multiplexed immunolabeling technique, and TACTIC, a single-cell-level, organ-scale image co-registration algorithm. 3D CYCLIC combines ultrafast, versatile 3D immunolabeling with a cleavable crosslinker that preserves signals by protecting bound antibodies during optical clearing while enabling their detachment for subsequent rounds of immunolabeling. TACTIC uses deep warping networks coupled with a propagation-based cell-pair search to co-register individual cells across whole-brain images acquired from multiple rounds of 3D CYCLIC labeling. 3D CYCLIC enabled 6-plex protein profiling of a mouse brain hemisphere, with images that can be combined with TACTIC for integrative analysis. 3D CYCLIC and TACTIC are versatile techniques and will therefore enable a holistic, unbiased understanding of diverse organs.

## Introduction

One of the most fundamental questions of biology is how the complexity of biological systems arises from a relatively simple genome^1,2^. Major drivers of this complexity are intermolecular and intercellular interactions^3^. Key insights into these interactions can be obtained by profiling biomolecules from individual cells with their cellular structures. Therefore, integrative imaging that can characterize molecular profiles and three-dimensional cellular structures can demystify how biological complexity emerges^4,5^.

Multiplexed immunofluorescence (mIF) techniques can label tens of target proteins from each biological specimen^6,7^. When combined with microscopy, mIF enables visualization of proteins along with cellular structures labeled by using immunolabeling or genetic approaches. mIF techniques can be classified by their multiplexing strategies, including fluorescence inactivation or antibody stripping coupled with cyclic immunolabeling (e.g., MELC^8^, MxIF^9^, SIMPLE^10^, 4i^11^), oligonucleotide conjugation of antibodies (e.g., CODEX^12^, SABER^13^), and spectral unmixing (e.g., PICASSO^14^, LUMos^15^). mIF has enabled precise cell-type identification using combinations of cell-type marker proteins^7,16^ and in-depth profiling of rare and highly heterogeneous tissues (e.g., clinical samples)^17,18^. However, current mIF techniques are limited to thin tissue sections, restricting their throughput and the range of cellular features — and thus intercellular interactions — they can identify. These limitations are particularly critical for organs with high regional heterogeneity and long-range three-dimensional intercellular interactions, such as the brain.

To achieve organ-scale mIF, antibody stripping coupled with a cyclic immunolabeling-based approach is the ideal multiplexing strategy, as it provides high signal intensity because it is the only multiplexing strategy free from molecular crowding, which decreases antibody binding and consequently lowers staining signals^6,19^. Signal intensity is particularly crucial for organ-scale immunostaining because its imaging noise and labeling background are higher than those of thin tissue imaging due to greater tissue light scattering^20^ and a larger pool of non-specific labeling^21^, respectively. However, realizing antibody stripping-based mIF in organ-scale tissues faces two major challenges. First, organ-scale immunolabeling requires an optical clearing process for imaging throughout organs or thick tissues; however, this process frequently causes dissociation of bound antibodies^22,23^. This issue can be prevented by crosslinking antibodies after labeling, which prevents antibody stripping^24^. Second, to make the organ-scale mIF meaningful, the information from images acquired across individual cyclic immunolabeling rounds should be integrated at the single-cell level. Existing techniques can co-register multiple whole-brain images with brain-region-level precision^25,26^ or align thin tissue-section (< 50 μm in thickness) images from cyclic immunolabeling with single-cell-level precision^27,28^. However, an algorithm capable of co-registering organ-scale images with single-cell-level precision is currently lacking.

In this study, we developed techniques that address the two challenges. We developed an organ-scale mIF technique compatible with off-the-shelf antibodies and conventional three-color fluorescence microscopy, termed 3D CYCLIC (three-dimensional **C**ellular protein profiling b**Y C**yclic **L**arge tissue **I**mmunolabeling using **C**leavable crosslinker), which achieves ultrafast, organ-scale, multiplexed immunolabeling using a cleavable crosslinker. Moreover, we developed TACTIC (**T**opology and **A**nchor-guided **C**ell-level **T**raining with **I**terative **C**o-registration), a framework that enables automatic single-cell-level co-registration of whole-brain images by integrating multiple rounds of images from 3D CYCLIC into a coherent, soma-indexed representation. 3D CYCLIC and TACTIC provide a powerful platform for single-cell-level protein profiling across whole organs, enabling unprecedented, system-wide views of complex biological processes.

## Results

### DTSSP enables both the protection of the immunostaining signal and the complete elution of bound antibodies

To reconcile the need to preserve the bound antibody during optical clearing with the need to elute them for antibody stripping-based cyclic immunostaining, we explored the use of a cleavable crosslinker. Among diverse crosslinkers, we found that 3,3’-dithiobis(sulfosuccinimidyl propionate) (DTSSP) protects bound antibodies, preserves immunofluorescence signals, and allows efficient antibody elution upon crosslinking cleavage. In contrast, paraformaldehyde (PFA), a widely used crosslinker for preserving bound antibodies during three-dimensional immunostaining^24,29^, does not allow complete antibody stripping (**Fig. 1a**).

**Figure 1.**
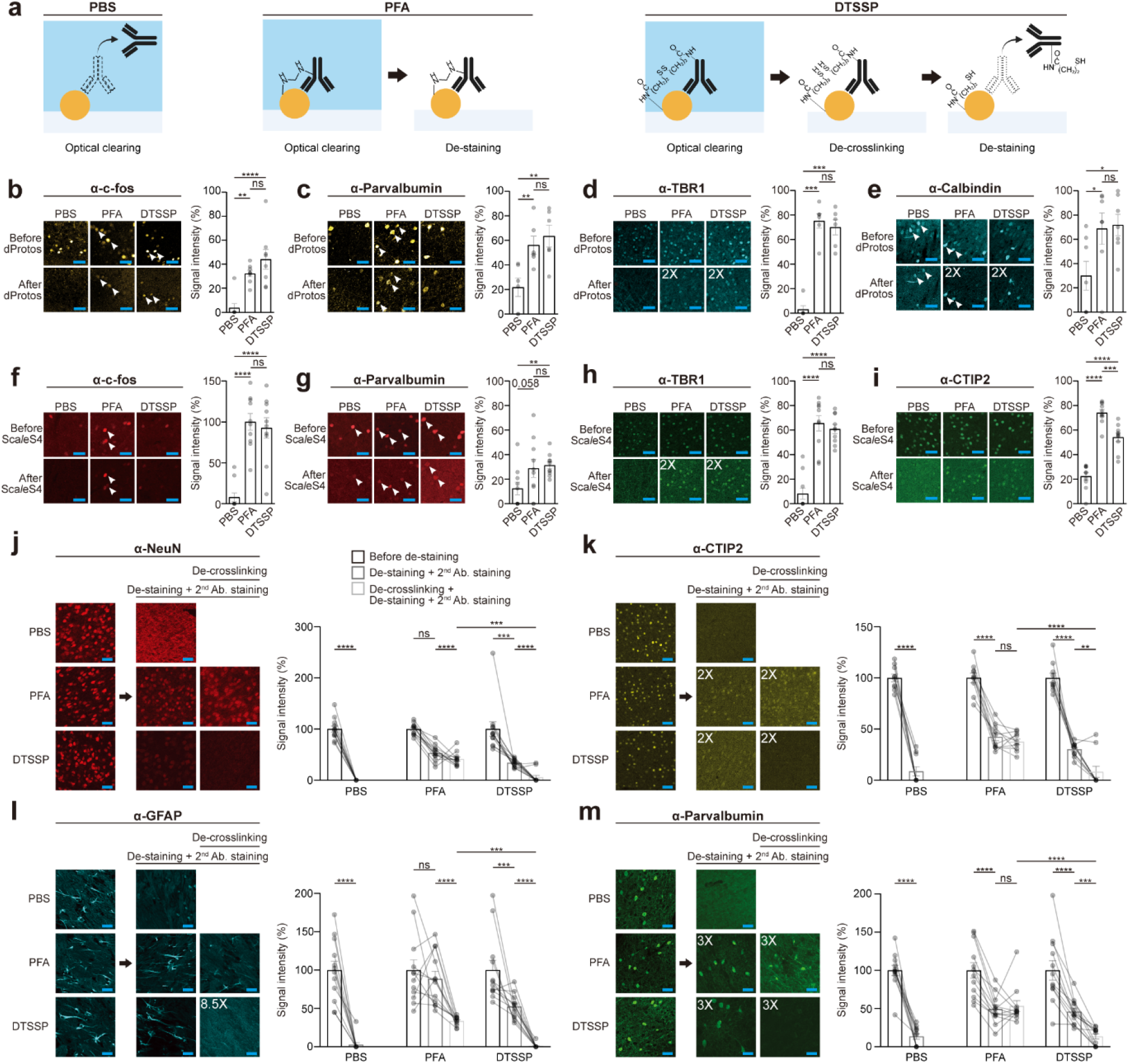
Effect of optical tissue clearing on antibody signal. **(a)**Schematic illustrating that optical clearing disrupts antigen-antibody binding, while DTSSP crosslinking is reversible, unlike PFA crosslinking. (**b-i**) Comparison of before and after optical tissue clearing of unfixed (PBS), PFA-fixed, and DTSSP-fixed tissues (*left*). Quantification of signal intensity (*right*). c-fos, *n* = 8. Parvalbumin, *n* = 6. NeuN, *n* = 10. Calbindin, *n* = 7. TBR1, *n* = 6. For antibodies, see **Supplementary Tables 1 and 2**. (2X = 2 times intensity gain by adjusting the display range of images). Images in **b-e** were optically cleared using dProtos, and images in **f-i** were cleared using Sca*l*eS4. Two-tailed paired *t*-test, \**P* < 0.05, \*\**P* < 0.01, \*\*\**P* < 0.001, \*\*\*\**P* < 0.0001; ns, not significant. (**j-m**) Immunolabeling signal before and after antibody de-staining of unfixed (PBS), PFA-fixed, and DTSSP-fixed tissues, with or without prior de-crosslinking or HIER (*left*). Quantification of signal intensity (*right*). TBR1, T-box brain transcription factor. CTIP2, Chicken ovalbumin upstream promoter transcription factor-interacting protein 2. GFAP, Glial fibrillary acidic protein. NeuN, *n* = 12. CTIP2, *n* = 10. GFAP, *n* = 12. PV, *n* = 12. “2X”, “3X” and “8.5X” indicate twofold, threefold and 8.5-fold increases, respectively, in displayed signal intensity by adjusting the image display range. Repeated-measures two-way ANOVA followed by Tukey’s multiple comparison test, \**P* < 0.05, \*\**P* < 0.01, \*\*\**P* < 0.001, \*\*\*\**P* < 0.0001; ns, not significant. Scale bars, 50 μm (blue). Error bars represent mean ± s.e.m. For used antibodies, see **Supplementary Table 2**.

To assess the effect of DTSSP crosslinking on the preservation of bound antibodies, we immunostained tissues with a panel of primary antibodies and treated them with either PFA or DTSSP. Tissues were then labeled with fluorophore-conjugated secondary antibodies, imaged, exposed to one of three optical clearing solutions^30-32^, and imaged again. By comparing tissue images before and after clearing, we found that all the optical clearing solutions substantially (∼62-97%) reduced immunofluorescence signals across all antibodies we tested (**Fig. 1b-i, Extended Data Fig. 1, Supplementary Table 1**). In contrast, DTSSP crosslinking after immunostaining preserved signals to a similar extent as PFA crosslinking (**Fig. 1b-i, Extended Data Fig. 1**). Signal loss upon optical clearing and signal preservation by DTSSP crosslinking were also observed for a dye (**Extended Data Fig. 2, Supplementary Table 2**). Together, these results indicate that DTSSP crosslinking protects immunofluorescence signals from optical clearing.

Next, we assessed the effect of DTSSP crosslinking and its cleavage on antibody elution. We treated immunolabeled and crosslinked tissues with an antibody elution protocol. When the tissues were re-stained with secondary antibodies to assess the remaining bound primary antibodies, we found that both PFA- and DTSSP-crosslinked tissues showed residual signals (**Fig. 1j-m**), confirming that crosslinking prevents antibody stripping. In contrast, when the DTSSP-crosslinked tissues were additionally treated with 1,4-dithiothreitol (DTT), an agent that cleaves DTSSP crosslinks^33^, we found that signals from the original primary antibodies were decreased (**Fig. 1j-m, Supplementary Table 1**; two-tailed *t*-test, \**P* < 0.05 or **** *P* < 0.0005) to levels not significantly different from zero (**Fig. 1j-m;** two-tailed *t*-test; anti-GFAP (Glial fibrillary acidic protein), *t*(11) = 1.00, *P* = 0.34; anti-NeuN, *t*(11) = 1.48, *P* = 0.17; anti-CTIP2 (Chicken ovalbumin upstream promoter transcription factor-interacting protein 2), *t*(9) = 1.49, *P* = 0.17); anti-PV (Parvalbumin), *t*(11) = 4.14, *P* = 0.0016). Complete elution could not be achieved in PFA-crosslinked tissues processed with Heat-Induced Epitope Retrieval (HIER), which is a process known to break PFA crosslinks using heat and high pH^34^ (**Fig. 1j-m;** two-tailed *t*-test; anti-GFAP, *t*(11) = 19.72, *P* = 6.2 × 10^-10^; anti-NeuN, *t*(11) = 10.89, *P* = 3.13 × 10^-7^; anti-CTIP2, *t*(9) = 13.36, *P* = 3.07 × 10^-7^; anti-PV, *t*(11) = 7.59, *P* = 1.07 × 10^-5^). This suggests that the cleavability of DTSSP enables complete elution of primary antibodies. To apply DTSSP crosslinking to organ-scale mIF, we developed a protocol that can crosslink bound antibodies throughout the mouse brain hemisphere (**Extended Data Fig. 3**). Collectively, these results indicate that DTSSP crosslinking can preserve immunolabeling signals during optical clearing and, after cleavage, enable complete detachment of primary antibodies throughout the mouse brain, thereby enabling subsequent cyclic immunolabeling of the same tissue.

### 3D CYCLIC enables mIF of the mouse brain hemisphere

Next, we integrated DTSSP-based reversible crosslinking with a modified version of eFLASH^35^, an ultrafast and versatile organ-scale immunolabeling technique, to enable organ-scale mIF (**Fig. 2a**). The resulting technique, termed 3D CYCLIC, begins with rapid primary antibody staining of delipidated mouse organs via a modified version of eFLASH, followed by DTSSP treatment and secondary antibody staining. The stained organ is then cleared with an optical clearing solution and imaged by light-sheet fluorescence microscopy. After imaging, the sample is incubated in a de-crosslinking solution (100 mM DTT in phosphate-buffered saline (PBS)) to cleave the DTSSP crosslinks, which enables antibody elution and the subsequent round of organ-scale immunolabeling (**Fig. 2a**).

**Figure 2.**
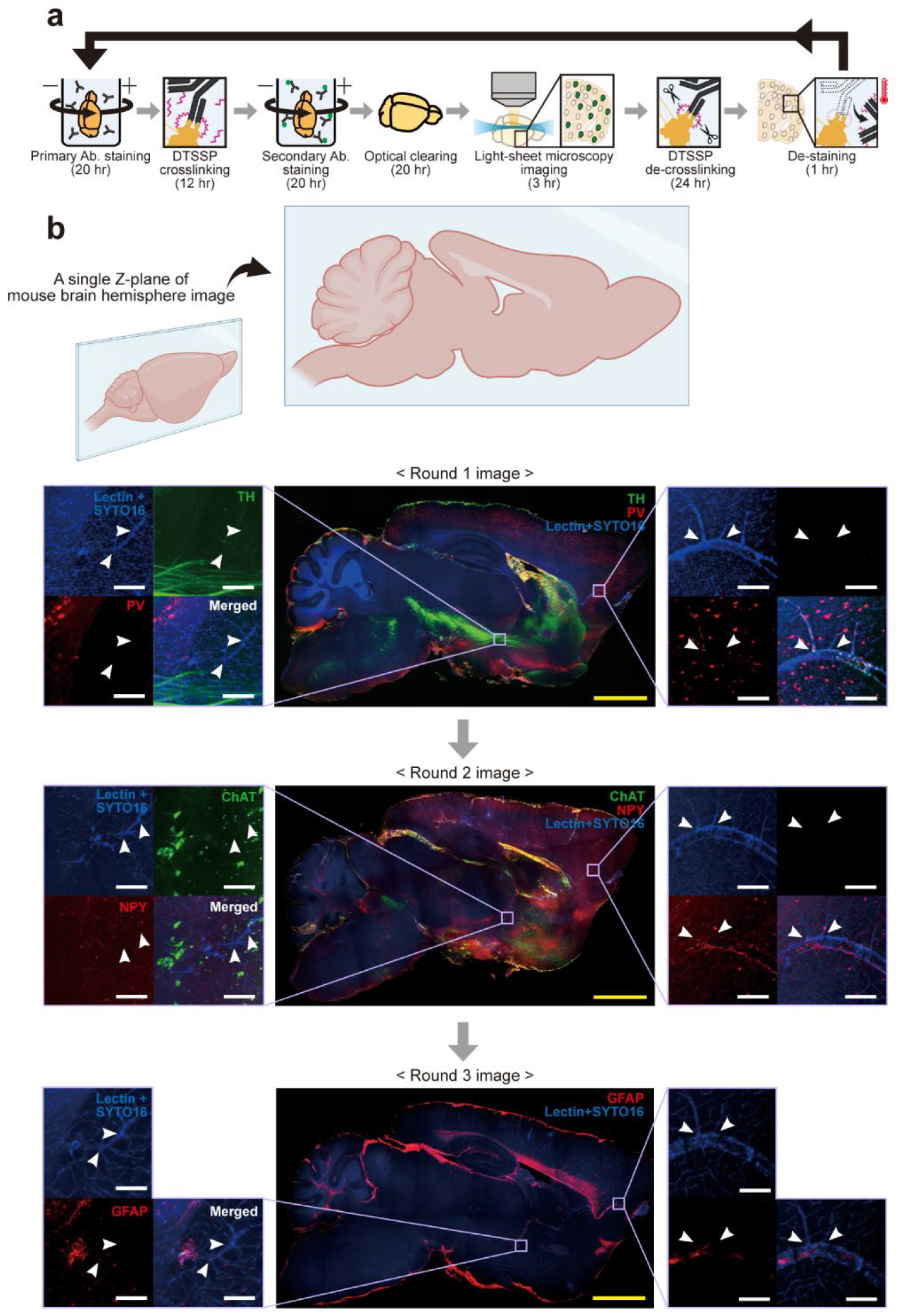
3D CYCLIC enables a multi-round molecular profile of the mouse hemisphere. (**a**) A schematic representation of the 3D CYCLIC workflow. (**b**) Representative single z-plane images of the mouse brain hemisphere from each staining round, with zoomed-in views of the indicated regions. Arrows indicate identical vascular landmarks across rounds. TH, Tyrosine hydroxylase; ChAT, Choline acetyltransferase; NPY, Neuropeptide-Y. Scale bars, 2 mm (yellow), 100 μm (white). For used antibodies, see **Supplementary Table 3**.

We applied 3D CYCLIC to a mouse brain hemisphere to demonstrate multiplexed, organ-scale profiling of six cell-type marker proteins. We included Lectin and SYTO16, a blood vessel marker and a nuclear dye, respectively, in each round of 3D CYCLIC as fiducial markers across individual round images. Using characteristic blood vessel structural landmarks, we confirmed that each staining round successfully labeled a distinct set of protein targets without interference from signals from previous rounds (**Fig. 2b, Supplementary Table 3**). These results demonstrate that 3D CYCLIC enables multiplexed, organ-scale protein profiling in intact organs.

### TACTIC enables single-cell-level co-registration of organ-scale mIF images

3D CYCLIC generates a series of organ-scale images capturing complementary molecular information from the same organ. If fiducial marker-expressing cells across individual round images throughout the organ can be co-registered, the molecular information from individual rounds can be integrated. This enables organ-scale, multiplexed protein profiling of individual fiducial marker-expressing target cells, including identification of their cell types and cellular states.

We enabled such image co-registration by developing TACTIC, which is a feedback-loop system comprising three modules: TACTIC-pair, -morph, and -propagate (**Fig. 3a**). The loop starts with seed-point cell pairs manually identified from each round of whole-brain images. Near the seed points, TACTIC-pair searches for candidate anchor cell pairs (*C*) with dense descriptors trained with morphology and topology embeddings. The anchor cell pairs let the TACTIC-morph module predict the local deformation field (*ϕ*) and conduct image warping through a spatial transformer network (STN)^36^. Around the finalized anchor cell pairs, TACTIC-propagate logs the iteration process and directs propagation toward maximum volume coverage while leveraging sustained pair-detection accuracy. If anchor cell pairs for propagation are found, the feedback loop continues to iterate. Otherwise, the process exits the loop, and TACTIC processing is finished.

**Figure 3.**
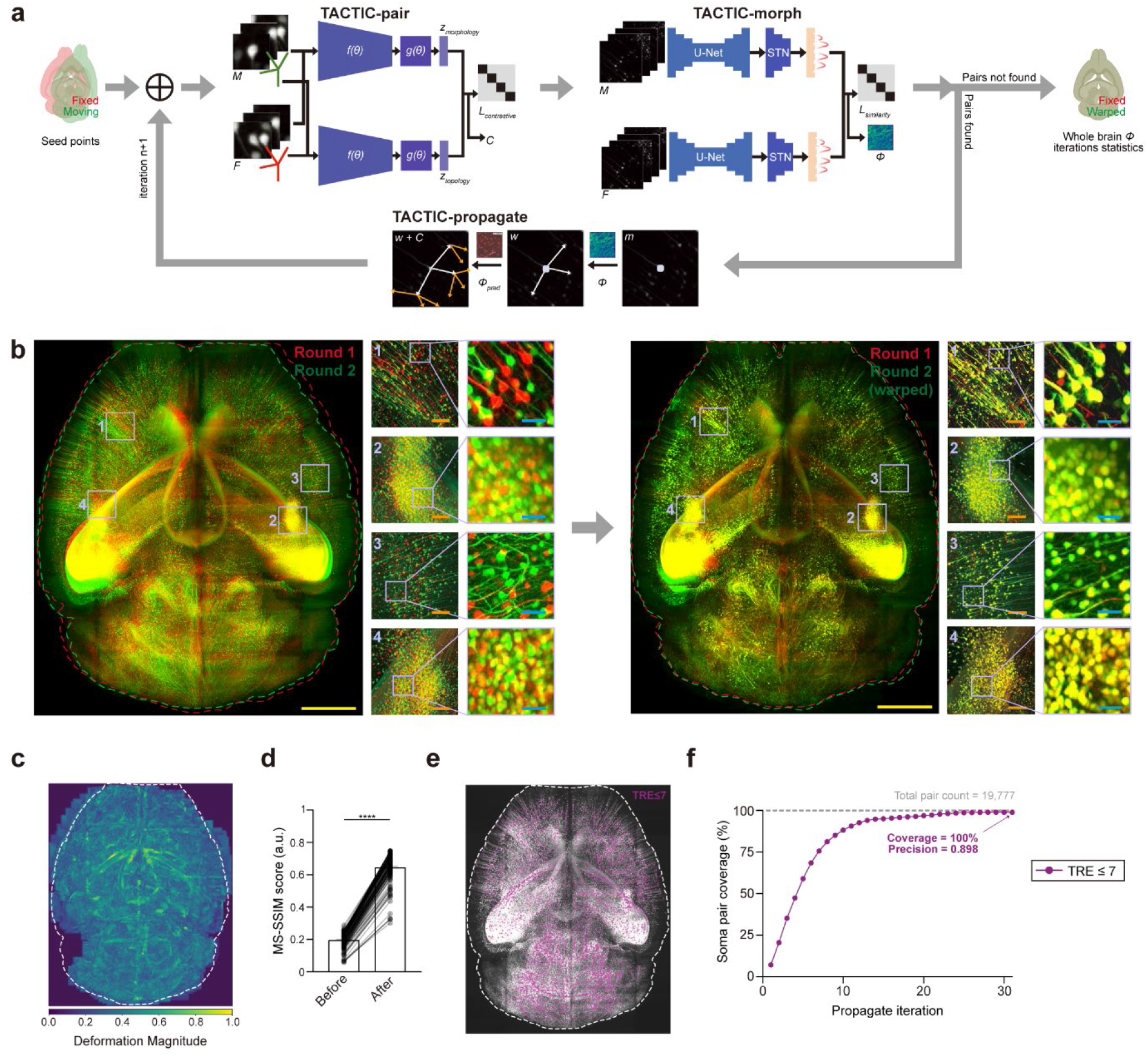
TACTIC enables single-cell-level co-registration of whole-brain images from a mouse. (**a**) Schematic of the TACTIC framework. *f(θ)*, feature encoder; *g(θ)*, feature projector; z_*morphology*_, morphology embedding; z_*topology*_, topology embedding; *L*_*contrastive*_, contrastive loss; *C*, candidate pairs; STN, Spatial Transformer Network; *ϕ*, deformation field; *ϕ*_*pred*_, deformation field with candidate pairs. (**b**) Maximum-intensity projection (MIP) images of a fixed brain image (green) and a moving brain image (red) before (*left*) and after (*right*) TACTIC co-registration. The fixed and moving brain images correspond to the image acquired before and after harsh treatment, respectively. Four representative brain regions are shown at two magnification levels to highlight the soma-level correspondence achieved after co-registration. (**c**) Whole brain MIP image of the magnitude of the predicted deformation field, ‖u‖, normalized to [0, 1] by the 99^th^ percentile across foreground voxels. Brighter hues indicate a larger deformation magnitude. (**d**) Comparison of MS-SSIM (Multi-scale Structural SIMilarity) scores between moving and fixed patches before and after TACTIC-morph-mediated warping. Higher MS-SSIM scores indicate greater structural similarity between the image pairs. Each scatter point on the bars represents the MS-SSIM of an individual patch. Warping increases similarity for all evaluated tiles. Two-tailed paired *t*-test, \*\*\*\**P* < 0.0001. (**e**) Registered eGFP+ neuron pairs overlaid on the whole-brain MIP image. Pairs with a target registration error (TRE) threshold of ≤ 7 voxels are regarded as co-registered. (**f**) Cumulative coverage of registered soma pairs as a function of TACTIC-propagate iterations. Coverage is expressed as a percentage of the total number of eGFP+ neuron pairs (19,777) with a TRE ≤ 7 voxels. Scale bars, 2 mm (yellow), 200 μm (orange), 50 μm (blue). Error bars represent mean ± s.e.m.

To test whether TACTIC can co-register two rounds of images generated from 3D CYCLIC with single-cell accuracy, we prepared two whole-brain images from the same Thy1-GFP-M mouse brain^37^. A whole-brain image was acquired after de-lipidation and optical clearing, and another whole-brain image was acquired from the same brain after a further harsh treatment corresponding to ∼10 rounds of whole-brain antibody stripping processes (80 °C, 4 hours of treatment in pH 9, 300 mM sodium dodecyl sulfate (SDS), 100 mM sodium sulfite, 10 mM boric acid solution). By applying TACTIC to the two whole-brain images using eGFP as a fiducial marker, we could co-register eGFP+ neurons of the two images at a single-cell resolution (**Fig. 3b**). The co-registration is enabled by 3D image warping, whose degree is heterogeneous across brain regions (**Fig. 3c, Extended Data Fig. 4**), which resulted in a significant increase in the similarity between the two images (**Fig. 3d**) and the co-registration of all eGFP+ neurons in the brain (19,777/19,777) with 89.8% precision (**Extended Data Fig. 5**). Across all target registration error (TRE) ranges (≤ 1, ≤ 3, ≤ 5, ≤ 7 voxels), matched soma pairs were found throughout the brain (**Extended Data Fig. 6**), indicating that TACTIC propagated and warped globally. These results suggest that TACTIC is an algorithm for comprehensive single-cell-level co-registration of whole-brain images, which might be applicable to individual round organ-scale images from the 3D CYCLIC labeling.

## Discussion

Intermolecular and intercellular interactions underlie complex biological functions and dysfunctions^3^. Single-cell omics techniques have advanced our understanding of intermolecular interactions by profiling molecules in individual cells^38,39^. The need to integrate rich molecular profiles with spatial information has led to the development of spatial omics techniques. Among those, spatial proteomic techniques^40-42^ are particularly powerful because they visualize proteins, the major biomolecules of biological function; however, they are intrinsically limited in their ability to comprehensively capture cellular morphology, which is critical for revealing intercellular interactions. mIF can inherently integrate protein profiles and cellular architecture at the single-cell level. By expanding mIF and its downstream analysis to the single-cell whole-organ level, 3D CYCLIC and TACTIC together have the potential to enable, for the first time, high-dimensional integrative mapping of protein profiles and cellular architectures across the entire mammalian brain.

This study achieved organ-scale mIF by addressing two key challenges: the hurdles of using a crosslinker and of integrating organ-scale images across multiple rounds. Organ-scale immunolabeling requires the use of a crosslinker because optical clearing processes, which are a prerequisite for organ-scale light microscopy^20,43^, frequently denature and detach bound antibodies from immunostained tissues, as their pH, osmolarity, and ionic strength deviate from those of antibody-antigen binding conditions^44,45^. Thus, crosslinkers such as PFA have been used to anchor bound antibodies^19,23^. However, this crosslinking prevents antibody stripping (**Fig. 1j-m**), thus making organ-scale mIF impossible. 3D CYCLIC overcomes this hurdle by using a cleavable crosslinker, DTSSP. For integrating images from multiple rounds, conventional 2D mIF techniques physically anchored tissue sections to glass slides and applied image co-registration algorithms^46,47^. However, this approach is not applicable to organ-scale tissues, because mounting on slide glasses or in cuvettes interferes with uniform immunolabeling, and the computational cost of image alignment scales quadratically with cell counts and with the magnitude of tissue deformation, making single-cell-level organ-scale image co-registration computationally unaffordable^48^. TACTIC resolves this challenge by combining deep warping networks with a propagation-based cell-pair search, achieving single-cell-level organ-scale image co-registration. Moreover, given the modest tissue deformation and preserved antigenicity observed across 3D CYCLIC rounds (**Extended Data Fig. 7**), these data suggest that additional rounds of 3D CYCLIC, combined with TACTIC, could further increase plex without compromising co-registration accuracy. To the best of our knowledge, 3D CYCLIC and TACTIC provide the first demonstration of organ-scale multiplexed immunolabeling and single-cell, organ-scale image co-registration, respectively.

Organ-scale mIF enabled by 3D CYCLIC and TACTIC can have diverse applications. 3D CYCLIC is compatible with diverse organs and antibodies, and, when combined with TACTIC, can enable unbiased, sensitive profiling of cell-type subtypes and cellular states^49,50^. Beyond cell phenotyping, the organ-scale mIF can profile 3D tissue architectures, including tumor microenvironments, follicular structures in lymphoid organs, and blood and lymphatic vessels^51^. These analyses are especially useful for biological samples with high inter-sample heterogeneity, such as clinical samples^52,53^. Furthermore, by being optimized for co-registering diverse cellular, subcellular, or anatomical fiducial markers throughout organs, TACTIC can enable integration of diverse imaging modalities, such as integrative imaging of neural activity of a neuron (acquired using *in vivo* two-photon calcium imaging) with its whole-brain projection pattern and downstream connected neuron’s cell type profiles (acquired using *ex vivo* 3D histology).

In summary, by developing 3D CYCLIC and TACTIC, we enabled organ-scale mIF. Based on their versatility—being compatible with diverse off-the-shelf antibodies and applicable to multiple organs—these two techniques will galvanize efforts to understand how biological functions and dysfunctions arise from intercellular and intermolecular interactions.

## Data Availability

All raw data described in this study are available from the corresponding authors upon request.

## Contributions

S.K., W.C., and Y.-G.P. developed 3D CYCLIC. H.P. and Y.-G.P. developed TACTIC. S.K. and W.C. performed the majority of *ex vivo* experiments with the help of S.Y. and T.C.. S.K., S.Y., and C.E.P. analyzed images. S.K., W.C., H.P., and Y.-G.P. wrote the paper with input from all authors.

## Corresponding author

Young-Gyun Park, Ph.D.

Korea Advanced Institute of Science and Technology (KAIST),

Department of Bio and Brain Engineering,

Daejeon, Republic of Korea

## Acknowledgements

The authors thank the members of the NBML (Neurotechnique and Brain Mapping Laboratory) for helpful discussions.

## Extended Data

**Extended Data Fig. 1.**
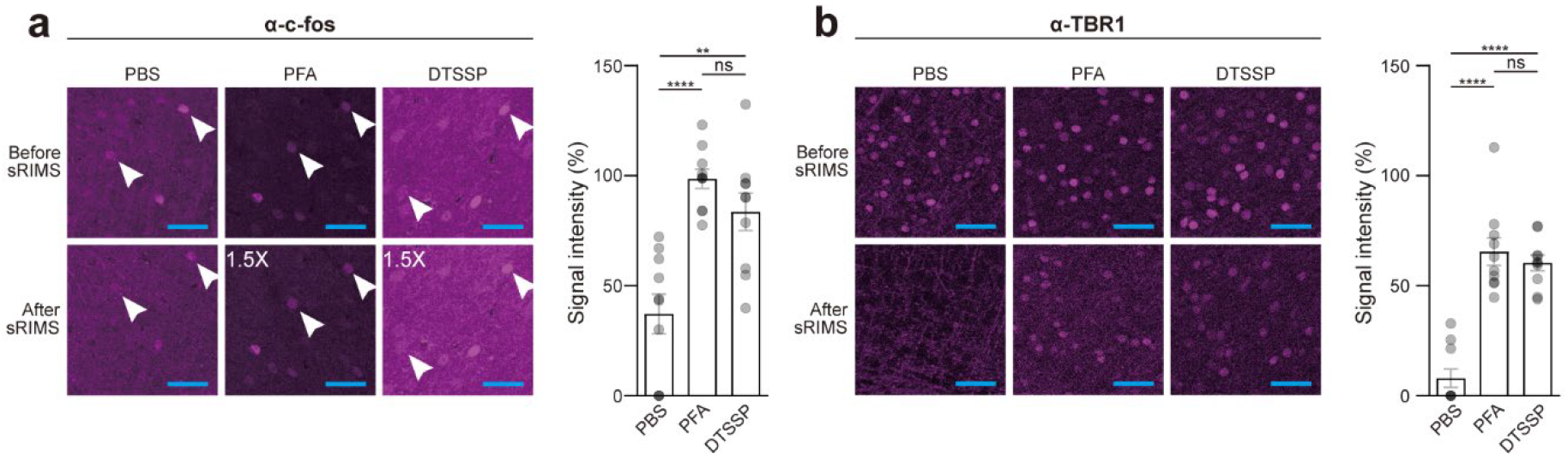
Effect of tissue optical clearing using sRIMS on antibody signal. (**a-b**) Comparison of immunostaining signal loss by sRIMS incubation in PFA-crosslinked (PFA), DTSSP-crosslinked (DTSSP), and non-crosslinked tissues (PBS). Representative images (*left*) and signal intensity quantification (*right*). c-fos, *n* = 10. TBR1 (T-box brain transcription factor), *n* = 10; “1.5X” indicates a 1.5-fold increase in displayed signal intensity by adjusting the image display range. Two-tailed paired *t*-test, \*\**P* < 0.01, \*\*\*\**P* < 0.0001; ns, not significant. Scale bar, 50 μm. Error bars represent mean ± s.e.m. For used antibodies, see **Supplementary Table 1**.

**Extended Data Fig. 2.**
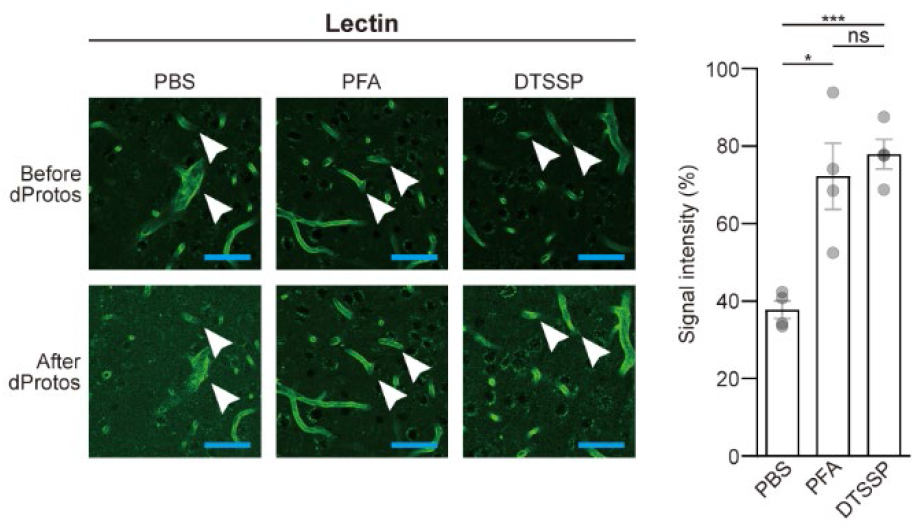
Effect of tissue optical clearing on dye signal. Comparison of lectin signal loss by dProtos incubation in PFA-crosslinked (PFA), DTSSP-crosslinked (DTSSP), and non-crosslinked tissues (PBS). Representative images (*left*) and signal intensity quantification (*right*). Lectin, *n* = 4. Two-tailed paired *t*-test, \**P* < 0.05, \*\*\**P* < 0.001; ns, not significant. Scale bar, 50 μm. Error bars represent mean ± s.e.m. For the used dye, see **Supplementary Table 1**.

**Extended Data Fig. 3.**
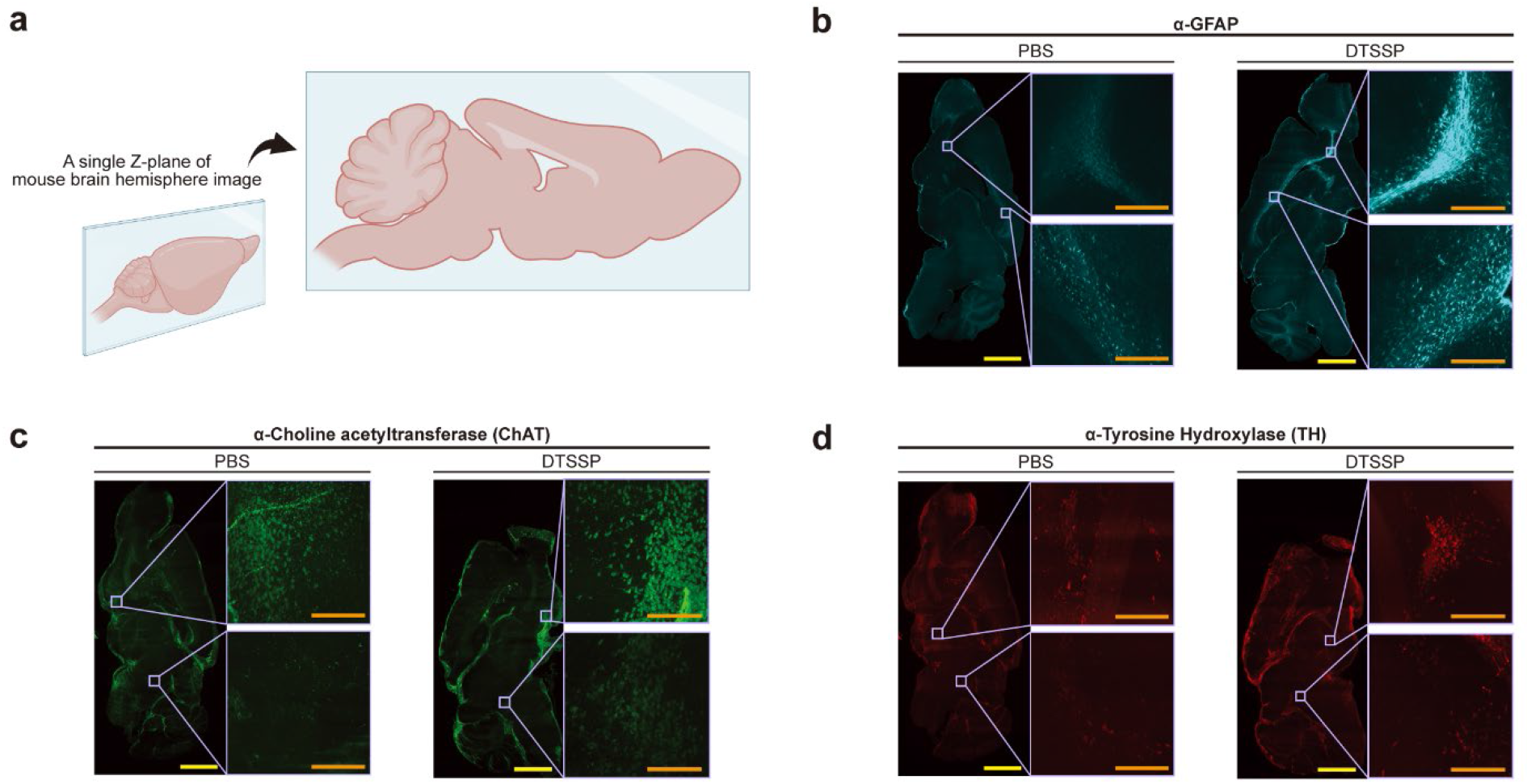
DTSSP enhances signal quality of organ-scale immunolabeling. (**a**) Schematic illustrating the experimental workflow for mouse hemisphere staining and 3D imaging. (**b-d**) Representative single z-plane images of mouse hemispheres. DTSSP-crosslinked hemispheres (DTSSP) showed higher signal intensity than non-crosslinked hemispheres (PBS). GFAP, Glial fibrillary acidic protein. Scale bars, 2 mm (yellow), 200 μm (orange).

**Extended Data Fig. 4.**
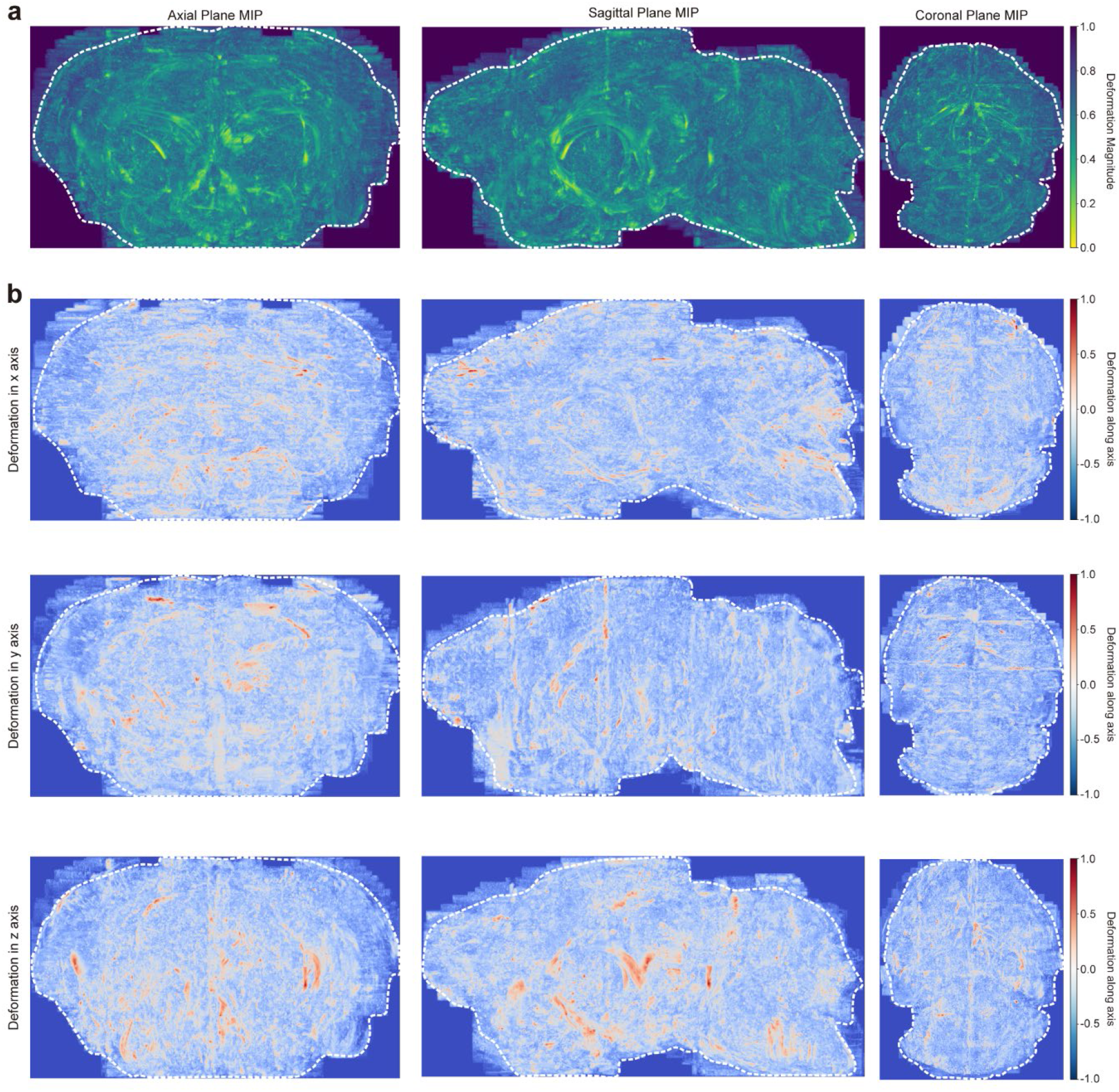
Brain-wide deformation field computed by the TACTIC-morph module. (**a-b**) Deformation field estimated during iterative TACTIC-propagate using the TACTIC-morph module. Maximum-intensity projection (MIP) images are shown in axial, sagittal, and coronal orientations (columns). (**a**) Deformation magnitude ‖u‖, normalized to [0, 1] by the 99^th^ percentile across foreground voxels. (**b**) Signed displacement components along the x-, y-, and z-axes (rows), jointly normalized to [−1, +1] by a single shared scale, preserving sign and enabling cross-axis comparison. MIP images emphasize voxels with the greatest deformation along the projection direction, enabling rapid identification of regions undergoing the greatest spatial adjustment during propagation.

**Extended Data Fig. 5.**
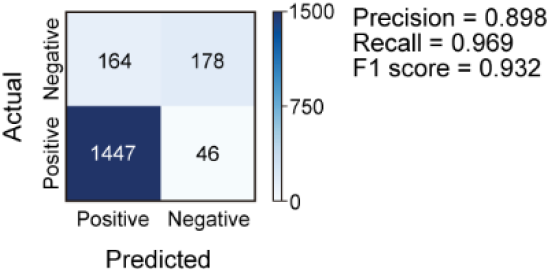
Manual validation of TACTIC-pair module predictions. Confusion matrix comparing TACTIC-pair calls (Predicted; x-axis) with manual annotations (Actual; y-axis) for TACTIC-pair module validation on held-out dataset. True positives (TP) and true negatives (TN) indicate correctly predicted soma pairs, whereas false positives (FP) and false negatives (FN) indicate incorrectly predicted paired and unpaired soma. Precision (TP / (TP + FP)) measures the fraction of predicted pairs that are correct, recall (TP / (TP + FN)) measures the fraction of true pairs that are recovered, and the F1 score is their harmonic mean (2TP / (2TP + FP + FN)).

**Extended Data Fig. 6.**
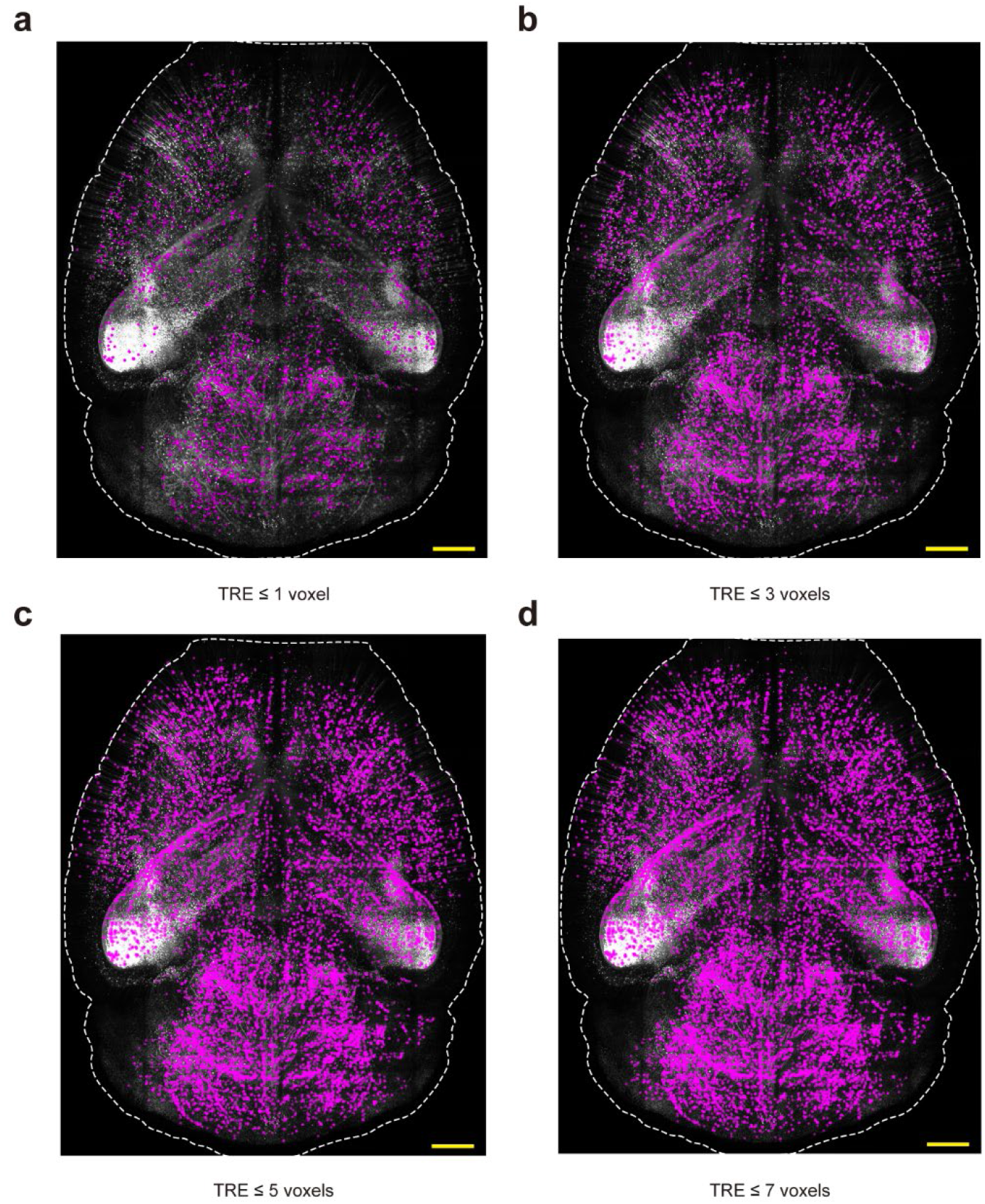
Spatial distribution of successfully registered soma pairs across increasing TRE thresholds. (**a-d**) Representative whole-brain MIP images showing soma-pair matches (magenta) that meet target registration error (TRE) cutoffs of ≤ 1, ≤ 3, ≤ 5, and ≤ 7 voxels. The grayscale image provides anatomical context, and the dashed white contour outlines the brain boundary. Scale bar, 2 mm.

**Extended Data Fig. 7.**
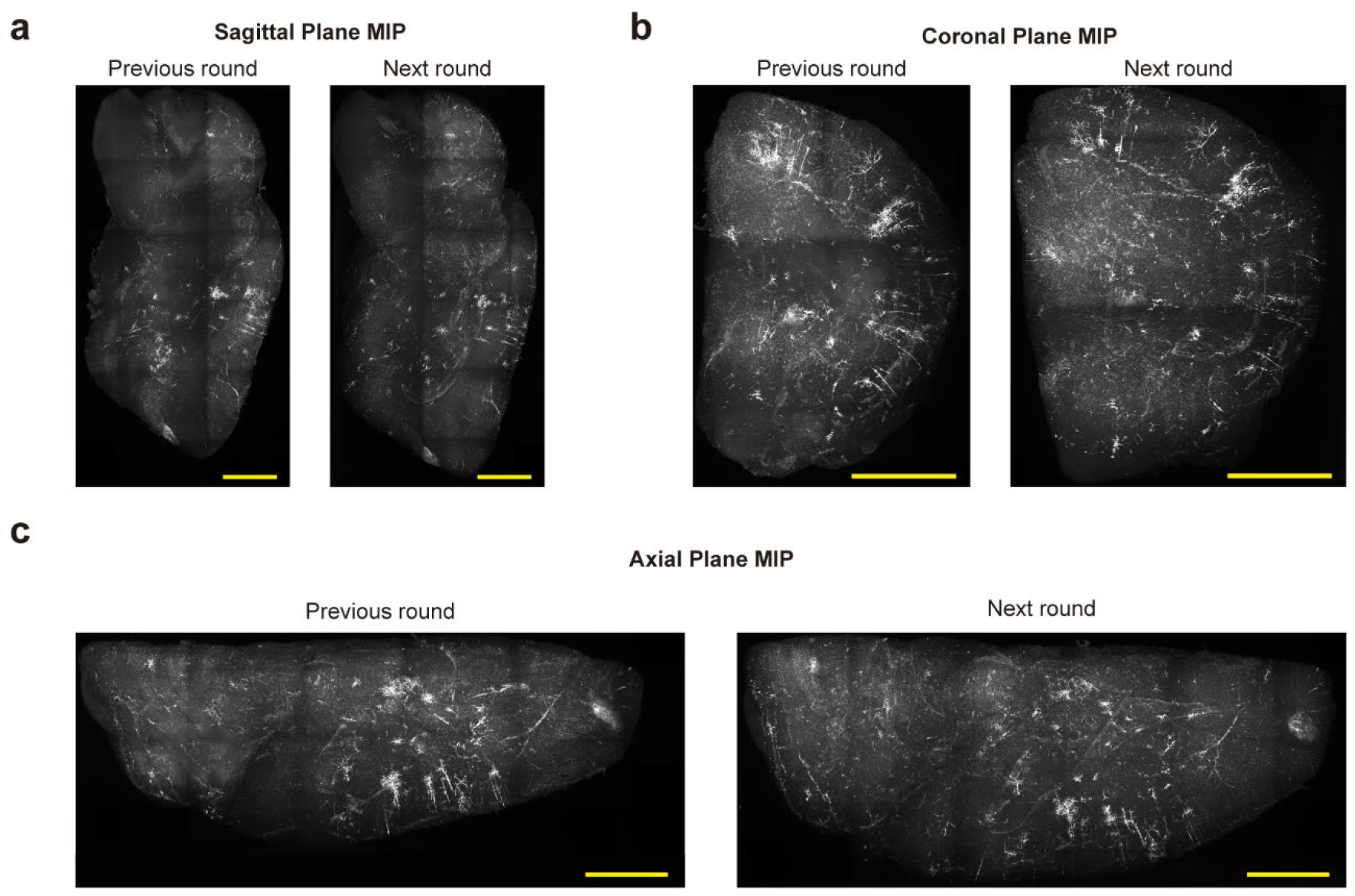
Whole-brain morphology shows no visible tissue deformation or loss of antigenicity across consecutive 3D CYCLIC rounds. (**a-c**) Two images of the same mouse brain hemisphere, acquired in consecutive immunolabeling rounds, are shown to assess brain deformation and signal quality across rounds. (**a**) Sagittal, (**b**) coronal, and (**c**) axial plane MIP images. Scale bar, 2 mm.

## Supplementary information

**Supplementary Table 1.**
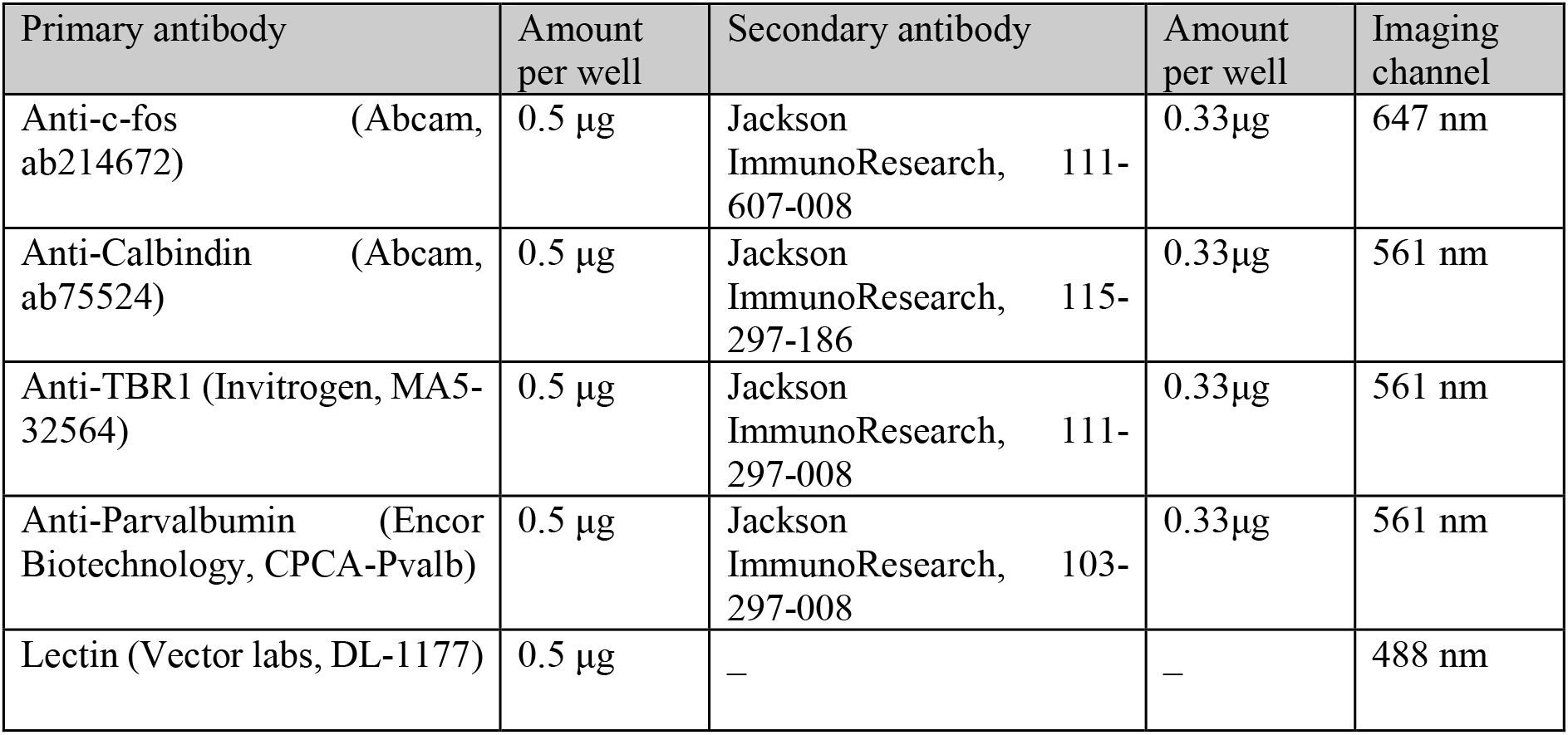
Antibodies and imaging conditions used for Figure 1b-e, Extended Data Fig. 1 and Extended Data Fig. 2.

| Primary antibody | Amount per well | Secondary antibody | Amount per well | Imaging channel |
| --- | --- | --- | --- | --- |
| Anti-c-fos (Abcam, ab214672) | 0.5 µg | Jackson ImmunoResearch, 111-607-008 | 0.33µg | 647 nm |
| Anti-Calbindin (Abcam, ab75524) | 0.5 µg | Jackson ImmunoResearch, 115-297-186 | 0.33µg | 561 nm |
| Anti-TBR1 (Invitrogen, MA5-32564) | 0.5 µg | Jackson ImmunoResearch, 111-297-008 | 0.33µg | 561 nm |
| Anti-Parvalbumin (Encor Biotechnology, CPCA-Pvalb) | 0.5 µg | Jackson ImmunoResearch, 103-297-008 | 0.33µg | 561 nm |
| Lectin (Vector labs, DL-1177) | 0.5 µg | — | — | 488 nm |

**Supplementary Table 2.** Antibodies and imaging conditions used for Figure 1j-m.

| Primary antibody | Amount per well | Secondary antibody | Amount per well | Imaging channel |
| --- | --- | --- | --- | --- |
| Anti-NeuN (CST, 12943S) | 2.5 μl | Jackson ImmunoResearch, 711-295-152 | 0.5 μg | 561 nm |
| Anti-CTIP2 (Abcam, ab18465) | 0.75 μg | Jackson ImmunoResearch, 712-545-153 | 0.5 μg | 488 nm |
| Anti-GFAP (CST, 3670S) | 1.67 μl | Jackson ImmunoResearch, 715-605-150 | 0.5 μg | 647 nm |
| Anti-PV (Invitrogen, PA1-933) | 0.5 μg | Jackson ImmunoResearch, 711-295-152 | 0.5 μg | 561 nm |

**Supplementary Table 3.**
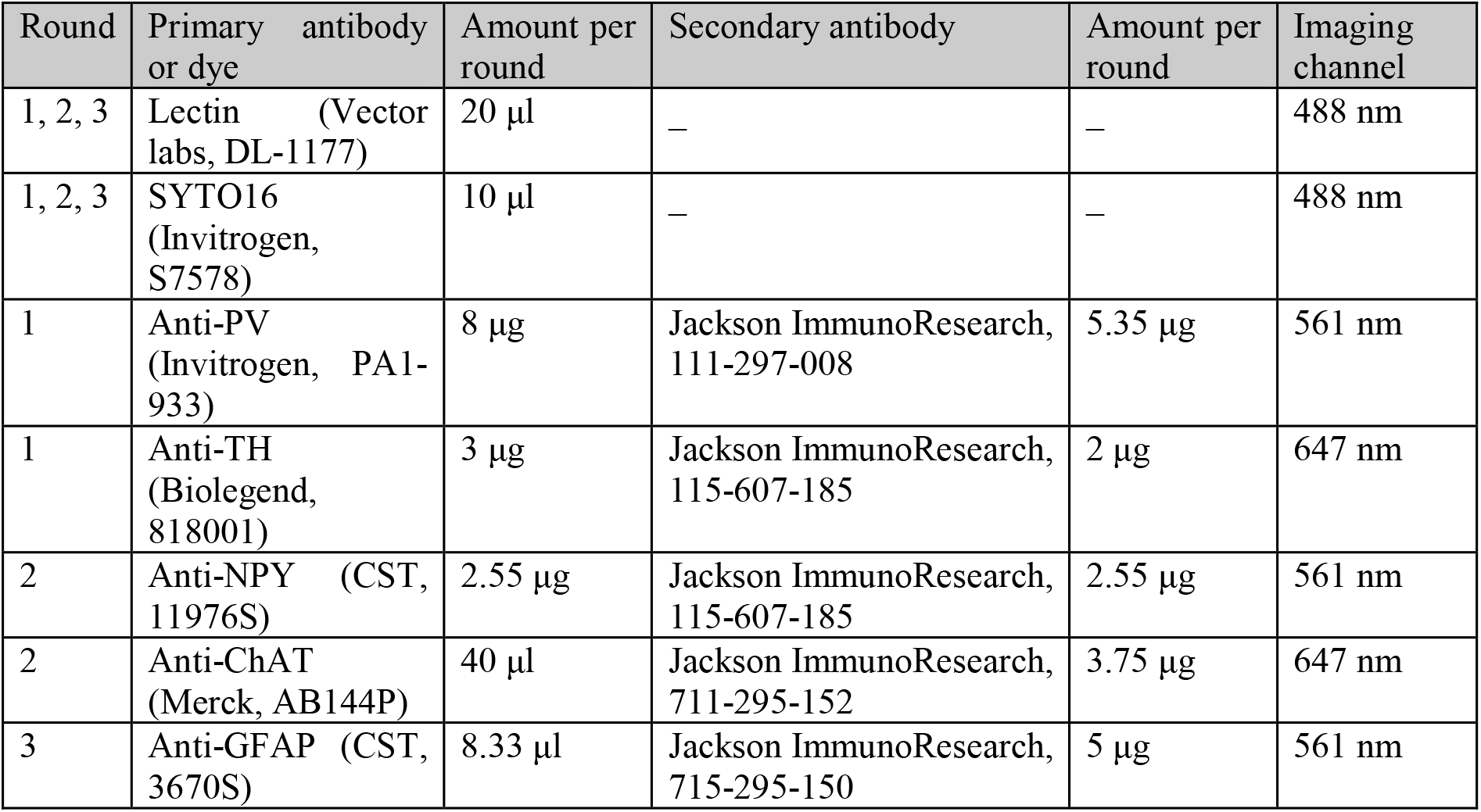
Antibodies and imaging conditions used for Figure 2.

| Round | Primary antibody or dye | Amount per round | Secondary antibody | Amount per round | Imaging channel |
| --- | --- | --- | --- | --- | --- |
| 1, 2, 3 | Lectin (Vector labs, DL-1177) | 20 μl | – | – | 488 nm |
| 1, 2, 3 | SYTO16 (Invitrogen, S7578) | 10 μl | – | – | 488 nm |
| 1 | Anti-PV (Invitrogen, PA1-933) | 8 μg | Jackson ImmunoResearch, 111-297-008 | 5.35 μg | 561 nm |
| 1 | Anti-TH (Biolegend, 818001) | 3 μg | Jackson ImmunoResearch, 115-607-185 | 2 μg | 647 nm |
| 2 | Anti-NPY (CST, 11976S) | 2.55 μg | Jackson ImmunoResearch, 115-607-185 | 2.55 μg | 561 nm |
| 2 | Anti-ChAT (Merck, AB144P) | 40 μl | Jackson ImmunoResearch, 711-295-152 | 3.75 μg | 647 nm |
| 3 | Anti-GFAP (CST, 3670S) | 8.33 μl | Jackson ImmunoResearch, 715-295-150 | 5 μg | 561 nm |

